# Evidence for an energetic trade-off model linking inflammaging and immunosenescence in the US Health and Retirement Study and UK Biobank

**DOI:** 10.1101/2025.10.17.682903

**Authors:** Jacob E. Aronoff, Maximilien Franck, Alan A. Cohen, Benjamin C. Trumble

## Abstract

Later life is characterized by the development of chronic inflammation, termed inflammaging, alongside changes in immune cell profiles, or immunosenescence. While these features contribute to health risk, they have also been interpreted as adaptive remodeling of the immune system in response to accumulating somatic damage. Here we consider a recently developed theoretical framework to understand these processes as interrelated: the Brain-Body Energy Conservation model of aging. This model views functional declines, such as immunosenescence, as part of an energy conserving response to the rising energy expenditure of inflammaging. This response promotes short term survival against somatic damage at the expense of future health risk. For example, naïve T cells, which enhance defense against future infections, decline with age. We find evidence consistent with this model in the US Health and Retirement Study (HRS) and UK Biobank (UKB). TNFR1, a key marker of inflammaging, mediated 10% and 5% of the age-related declines in naïve CD4T and CD8T cells respectively in the HRS (n = 8,261). Consistent with an impaired immune response to future infections, TNFR1 also mediated 16% of the age-related increased risk of hospitalization or death from COVID-19 in the UKB (n = 522 hospitalized or died, full sample n = 40,638). GDF15, which is produced in response to metabolic stress and has been found to induce immune tolerance in response to chronic inflammation, mediated 28% of the TNFR1-related COVID-19 health risk, as well as 38% of the age-related increased risk independent of TNFR1.

## Introduction

Later life is accompanied by complex changes in immune function that are implicated in health risk. There is a pronounced decline in adaptive immunity, which has been referred to as immunosenescence. This is reflected for example in the involution of the thymus and resulting decline in naïve T cells, which play an important role in defense against novel infectious exposures (Blackwell et al., 2016; Fülöp et al., 2018; Peters, Delhey, Nakagawa, Aulsebrook, & Verhulst, 2019; Shanley, Aw, Manley, & Palmer, 2009). Later life is also accompanied by the development of chronic, systemic, and sterile inflammation, termed inflammaging (Franceschi et al., 2000; Fülöp et al., 2018; Fülöp et al., 2021). While these changes are often interpreted as immune dysfunction that contributes to aging and disease, some have proposed that they at least partly reflect immune remodeling that “makes the best of a bad situation” (Fülöp et al., 2018; Fülöp, Larbi, Hirokawa, Cohen, & Witkowski, 2020).

A new theoretical model provides a framework to understand the relationship between inflammaging and immunosenescence from the perspective of adaptive remodeling: the Brain-Body Energy Conservation (BEC) model of aging (Shaulson, Cohen, & Picard, 2024). The BEC model highlights the energetic cost of inflammaging, which is progressively activated in response to accumulating somatic damage. However, total body energy expenditure in late life does not increase (Pontzer et al., 2021). The suite of functional declines and tissue atrophy observed in later life, including immunosenescence, are therefore viewed as energy conserving responses.

Current evidence suggests inflammaging is energetically costly. For example, cellular injury over time can lead to the development and accumulation of senescent cells throughout the body. These cells have arrested cell division, yet they are continually active. Their pro-inflammatory signaling, termed the senescence-associated secretory profile (SASP) (Coppé et al., 2008), is a major contributor to inflammaging. These cells display markers of increased energy expenditure, such as greater mitochondrial density (Shaulson et al., 2024).

The BEC model draws from evolutionary life history theory (LHT), which seeks to understand how organisms allocate their finite resources, including energy, between competing functions to maximize survival and reproduction (Roff, 2002; Stearns, 1992). Central to LHT is the concept of trade-offs, as energy invested in one function comes at the expense of another. These trade-offs are often studied between three broad categories of growth, reproduction, and maintenance, with immune function considered part of maintenance (Clancy et al., 2013; Lochmiller & Deerenberg, 2000; McDade, 2003; Muehlenbein, Hirschtick, Bonner, & Swartz, 2010; Urlacher et al., 2018; Zuk & Stoehr, 2002). However, there is also evidence for potential trade-offs occurring within different components of the immune system, as each comes with different costs and benefits (McDade, Georgiev, & Kuzawa, 2016). Innate immunity provides non-specific and fast-acting defense against cellular damage, and its activation is a central component of inflammaging. In contrast, adaptive immunity primarily provides enhanced future protection against infections. This occurs through the proliferation and development of naïve T cells, which can respond to novel pathogens and develop memory for improved future defense (McDade et al., 2016; Paul, 2013). As a result, much of adaptive immunity provides little immediate survival benefit, suggesting it would be one of the first functions to be divested from with accumulating energy expenditure from inflammaging. Furthermore, rate of encounters with novel pathogens should decrease with aging, as more and more of the most common pathogens have already been encountered with increasing age; considering the increasing risk of autoimmunity as well, there may be substantial pressure to reduce adaptive immune investment with age.

Here we test the BEC model linking inflammaging to immunosenescence using data from two large human cohorts: The US Health and Retirement Study (HRS) and the UK Biobank (UKB). The HRS is a nationally representative study of US older adults, which provides publicly available data including several thousand individuals with measured cytokines and naïve T cells. We used naïve T cell counts as a proxy for investment in the immune repertoire, which decline with thymus involution in later life. We also used the cytokines interleukin (IL)-6 and soluble tumor necrosis factor receptor 1 (TNFR1), which are important orchestrators of inflammaging and are part of the SASP signaling (Franck, Tanner, et al., 2025; Minciullo et al., 2016; Partridge, Fuentealba, & Kennedy, 2020). We assessed whether IL-6 and TNFR1 mediated inverse associations between age and naïve T cells. We also assessed whether these cytokines mediated inverse associations between a collection of chronic diseases reflecting accumulated somatic damage and inflammaging and naïve T cells. This included hypertension or cardiovascular disease (CVD), diabetes, and cancer.

An expected cost of the trade-off between inflammaging and immunosenescence is impaired defense against novel infections. To assess this possibility, we relied on the UKB, which contains cytokine measures in a large sample (approximately 54,000 individuals) at baseline (years 2006-2010) as well as tracking of COVID-19 hospitalizations or deaths (years 2020-2024). This natural experiment allowed us to test whether greater inflammaging prospectively predicted greater vulnerability to a novel infection in the future. The proteomics data from the UKB contains IL-6 and TNFR1 to directly compare with the HRS as well as other important orchestrators of inflammaging: IL-1β and TNF-α. We assessed whether these cytokines mediated positive associations between age and hospitalization or death from COVID-19 over follow up. Similar to the HRS study, we also assessed whether these cytokines mediated positive associations between chronic diseases and hospitalization or death.

Others have noted previously that inflammation “activates many components of dormancy programs typically induced by nutrient scarcity” (A. Wang, Luan, & Medzhitov, 2019). An important mechanism in this process is likely GDF15, which signals metabolic stress and has been proposed as an important orchestrator of energy conservation (Shaulson et al., 2024). GDF15 plays a role in immune suppression or tolerance in response to chronic inflammation, which could increase vulnerability to novel viral infections like COVID-19 (Ahmed et al., 2022; Bu et al., 2024; Luan et al., 2019; Pence, 2022; Salminen, 2025). Since GDF15 was measured as part of the proteomics panel in the UKB, we assessed whether it mediated positive associations between pro-inflammatory cytokines and hospitalization or death from COVID-19.

Finally, while increased pro-inflammatory signaling has been observed with age, increased anti-inflammatory signaling has also been observed across multiple diverse human populations (Aronoff et al., 2025; Morrisette-Thomas et al., 2014). There are multiple possible reasons for the development of what has been termed anti-inflammaging (Franceschi et al., 2007; Minciullo et al., 2016). Although anti-inflammatory signaling is generally considered protective for health, there is also evidence that it reinforces inflammaging-driven immunosenescence. An experimental study in mice found that the anti-inflammatory cytokine IL-10 increased systemically in response to increasing IL-6 with age and was mechanistically involved in suppressing the immune response (Almanan et al., 2020). In addition, IL-10 has been found to exert its anti-inflammatory effects through metabolic reprogramming of macrophages (Ip, Hoshi, Shouval, Snapper, & Medzhitov, 2017), fitting with an energy conservation framework. We therefore considered whether IL-10 mediated positive associations between pro-inflammatory cytokines and risk of hospitalization or death from COVID-19 in the UKB.

**Figure 1.**
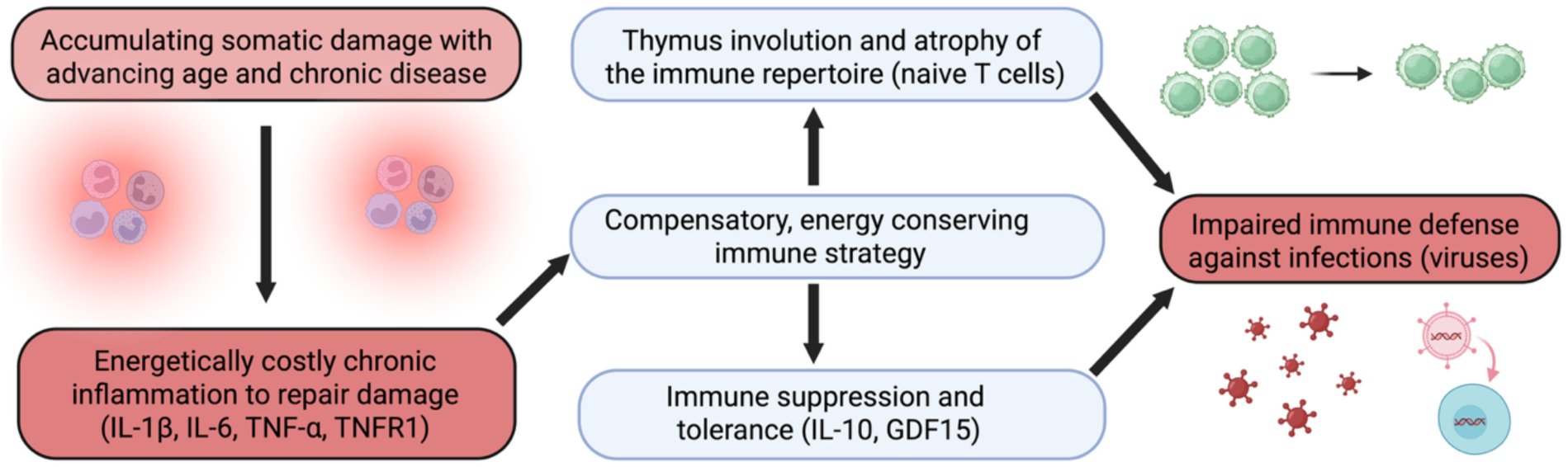
Conceptual figure of the study. Pro- and anti-inflammatory signaling drive energy-conserving immunosuppression in aging.

## Methods

### HRS sample

The HRS is a nationally representative study of US older adults. Here we focus on the 2016 cross-section that included a venous blood draw. Participants self-reported whether they had received a diagnosis of hypertension, a heart condition (cardiovascular disease), diabetes, or cancer (excluding skin). Measurement of cytokines and naïve T cells have been described previously (Crimmins, Faul, Thyagarajan, & Weir, 2017; Thyagarajan et al., 2022). Briefly, IL-6, IL-10, and TNFR1 were measured in serum with the Simple Plex assays on the ELLA System (Crimmins et al., 2017). Naïve T cells were measured using flow cytometry from whole blood samples collected in participants’ homes. Blood samples were shipped to the University of Minnesota and processed within 48 hours of collection. Samples were centrifuged to obtain peripheral blood mononuclear cells (PBMCs), which were then cryopreserved in liquid nitrogen. Flow cytometry was performed on thawed PBMC samples using either an LSRII flow cytometer or a Fortessa X20 instrument (BD Biosciences, San Diego, CA). Immunophenotyping data were analyzed using OpenCyto and FlowAnnotator. Counts of naïve T cells per milliliter of peripheral blood were obtained by multiplying the lymphocyte count by the percentage naïve T cells. Lymphocyte counts were obtained from a differential white blood cell count in an EDTA blood sample collected simultaneously and a Sysmex XE-2100 instrument (Sysmex America, Inc., Lincolnshire, IL) (Hunter-Schlichting et al., 2020; Thyagarajan et al., 2018; Thyagarajan et al., 2022).

### UKB sample

The UKB is a large population-based study of health that began with the recruitment of approximately 500,000 participants across the UK from 2006-2010. Participants self-reported whether they had hypertension, cardiovascular disease, diabetes, or cancer. Cytokines were measured in a subsample through proteomic profiling of blood plasma samples with the antibody-based Olink Explore 3072 PEA (Sun et al., 2023). Hospitalization or death from COVID-19 from January 2020 through April 2024 was recorded using hospital and death records. We specifically focused on cases in which COVID-19 was recorded as the primary diagnosis or cause of death.

### Analysis

To rule out potential confounding by acute infection, we excluded individuals with leukocytosis in both cohorts, using >11,000 leukocytes per microliter (µL) of blood as a cutoff. This excluded 2.87% of the HRS sample and 2.37% of the UKB. In addition, UKB participants who had died prior to January 1, 2020 (6.9%) or who were lost to follow-up (0.3%) were excluded from the analysis. This results in final sample sizes of n=8,261 for the HRS and n = 40,638 for the UKB. Within the UKB sample, n = 522 were either hospitalized or died from COVID-19 (n = 486 hospitalizations, n = 168 deaths).

In the HRS, cytokines were natural log-transformed due to positive skew. Naïve CD4T and CD8T cell counts were also positively skewed and natural log-transformed after a positive offset (+1) to adjust for zero values. UKB cytokine measures were approximately normally distributed and therefore not log-transformed. Cytokines in both cohorts, as well as naïve T cells, were then standardized (mean = 0, SD = 1), and outliers were winsorized to the 1^st^ and 99^th^ percentiles of the distribution.

We used ordinary least squares (OLS) regression models with the HRS venous blood sample 2016 weights to test associations between age, diseases, cytokines, and naïve T cells. In the UKB, logistic regression models tested associations between age, diseases, cytokines, and hospitalization or death from COVID-19. For cytokines showing results consistent with mediation, we conducted formal mediation analysis using a quasi-Bayesian approach with 1,000 simulations (Imai, Keele, & Tingley, 2010; Tingley, Yamamoto, Hirose, Keele, & Imai, 2014). The code used in the analysis can be found at: https://github.com/jakearonoff/hrs_ukb_immuneaging_tradeoff.

## Results

Descriptive statistics for the two cohorts are shown in **Table 1**. The UKB sample was substantially larger, while the HRS sample was older and had a higher prevalence of disease at the time of cytokine measurement. In both cohorts, there were slightly more women than men, with the difference being slightly greater in the HRS. All cytokines were positively correlated across both the HRS and UKB (**Supplementary Tables S1-2**). In the HRS, both IL-6 and TNFR1 were positively associated with age and disease (**Supplementary Table S3**). In the UKB, all pro-inflammatory cytokines were positively associated with age. Nearly all of these cytokines in the UKB were also positively associated with diseases, with the exception of IL-1β, which was not associated with diabetes or cancer (**Supplementary Table S4**).

**Table 1.**
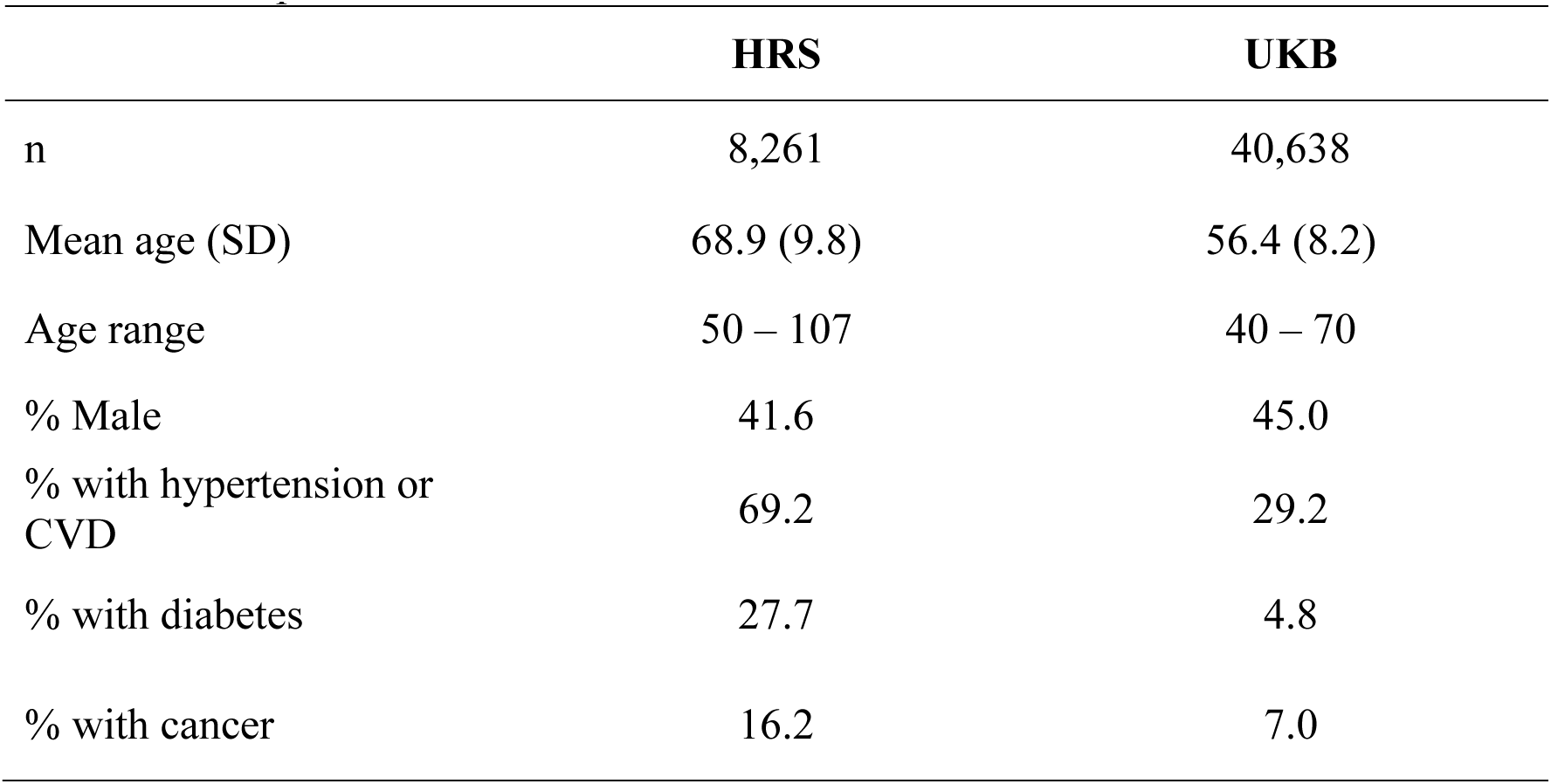
Descriptive statistics.

|  | HRS | UKB |
| --- | --- | --- |
| n | 8,261 | 40,638 |
| Mean age (SD) | 68.9 (9.8) | 56.4 (8.2) |
| Age range | 50 – 107 | 40 – 70 |
| % Male | 41.6 | 45.0 |
| % with hypertension or CVD | 69.2 | 29.2 |
| % with diabetes | 27.7 | 4.8 |
| % with cancer | 16.2 | 7.0 |

Regression models showing associations between age, diseases, cytokines, and naïve T cells in the HRS are shown in **Table 2**. In the base model predicting IL-10, one year of age was associated with a 0.012 standard deviation increase, while having hypertension or CVD, diabetes, and cancer were each associated with a 0.181, 0.231, and 0.112 increase respectively. Male sex was associated with a 0.098 standard deviation increase in IL-10. Consistent with mediation, age and disease associations were attenuated after adding IL-6 and TNFR1 to the model, while one standard deviation increase in IL-6 and TNFR1 was associated with a 0.184 and 0.366 standard deviation increase in IL-10 respectively. In the base model predicting naïve CD4T cells, a one-year increase in age was associated with a 0.018 standard deviation decrease in naïve CD4T cell counts. In addition, having hypertension or CVD, diabetes, and cancer were each associated with a 0.053, 0.115, and 0.270 standard deviation decrease respectively in naïve CD4T cells. Male sex was also associated with a 0.358 standard deviation decrease in naïve CD4T cells. Consistent with mediation, the addition of IL-6 and TNFR1 attenuated the negative associations with age and diseases, while one standard deviation increase in these cytokines was associated with a 0.034 and 0.061 standard deviation decrease in naïve CD4T cells respectively. In the final model with all predictors, VIF’s were the following: age = 1.25, sex = 1.01, hypertension or CVD = 1.17, diabetes = 1.10, cancer = 1.03, IL-6 = 1.25, TNFR1 = 1.49.

**Table 2.** Regression models predicting Naïve T cells in the US HRS. Cytokines and T cell counts were standardized, with resulting regression coefficients reflecting changes in standard deviations (n = 8,261). Estimates are reported as β (SE).

|  | IL-10 | IL-10 | CD4T | CD4T | CD8T | CD8T |
| --- | --- | --- | --- | --- | --- | --- |
| Age<br>(years) | 0.012***<br>(0.001) | -0.003*<br>(0.001) | -0.018***<br>(0.001) | -0.015***<br>(0.001) | -0.036***<br>(0.001) | -0.034***<br>(0.001) |
| Male<br>(0,1) | 0.098***<br>(0.020) | 0.098***<br>(0.018) | -0.358***<br>(0.021) | -0.358***<br>(0.021) | -0.308***<br>(0.017) | -0.310***<br>(0.017) |
| Hypertension<br>or CVD<br>(0,1) | 0.181***<br>(0.022) | 0.026<br>(0.020) | -0.053*<br>(0.023) | -0.027<br>(0.024) | -0.014<br>(0.019) | 0.003<br>(0.020) |
| Diabetes<br>(0,1) | 0.231***<br>(0.023) | 0.070**<br>(0.022) | -0.115***<br>(0.025) | -0.087***<br>(0.025) | -0.008<br>(0.021) | 0.011<br>(0.021) |
| Cancer<br>(0,1) | 0.112***<br>(0.027) | 0.047<br>(0.024) | -0.270***<br>(0.029) | -0.259***<br>(0.029) | -0.067**<br>(0.024) | -0.059*<br>(0.024) |
| IL-6 |  | 0.184***<br>(0.011) |  | -0.034**<br>(0.013) |  | 0.006<br>(0.011) |
| TNFR1 |  | 0.366***<br>(0.012) |  | -0.061***<br>(0.015) |  | -0.064***<br>(0.012) |

\* p < 0.05; \*\* p < 0.01; \*\*\* p < 0.001

Results for naïve CD8T cells were largely similar to CD4T, although overall slightly weaker. TNFR1 was inversely associated with naïve CD8T cells, and consistent with mediation its addition to the model attenuated the inverse age association. A similar pattern of results was observed for cancer, suggesting TNFR1 mediated the inverse association with naïve CD8T cells. However, IL-6, as well as diseases of hypertension or CVD and diabetes, were not associated.

A potential confounder in the HRS results is cytomegalovirus (CMV), which has been found to contribute to both inflammaging and immunosenescence (McElhaney et al., 2012). Therefore, as a sensitivity analysis, we considered additional models that included IgG antibodies against CMV. These models produced largely similar results (**Supplementary Table S5**). The inverse associations between TNFR1 and naïve T cells were nearly identical, while the inverse association between IL-6 and CD4T cells was slightly attenuated.

Mediation results for cytokines and naïve T cells in the HRS are shown in **Supplementary Table S6** and **Figure 2**. TNFR1 mediated 100% of the positive association between age and IL-10, as well as 10.1% and 5.0% of the inverse associations with naïve CD4T and CD8T cells respectively. IL-6 did not mediate any of these associations independently of TNFR1. For hypertension or CVD, TNFR1 mediated 74.2% of the positive association with IL-10 independent of IL-6, while IL-6 mediated 47.9% of the association independent of TNFR1. Further, TNFR1 mediated 30.7% of the negative association between hypertension or CVD and naïve CD4T cells independent of IL-6, while IL-6 mediated 12.1% of the association independent of TNFR1, although statistical significance for these mediation paths was weaker due to the slight overall association (p = 0.084 for TNFR1, p = 0.180 for IL-6). TNFR1 mediated 56.5% of the positive association between diabetes and IL-10 independently of IL-6, while IL-6 mediated 20.8% of the association independently of TNFR1. In addition, TNFR1 mediated 14.9% of the inverse association between diabetes and naïve CD4T cells independently of IL-6, while IL-6 mediated 3.6% of the association independently of TNFR1. For cancer, TNFR1 mediated 43.2% of the positive association with IL-10, while there was slight evidence that IL-6 also mediated 13.2% of the association independently of TNFR1 (p = 0.130). Finally, TNFR1 mediated 2.3% and 9.6% of the inverse associations between cancer and naïve CD4T and CD8T cells respectively.

**Figure 2.**
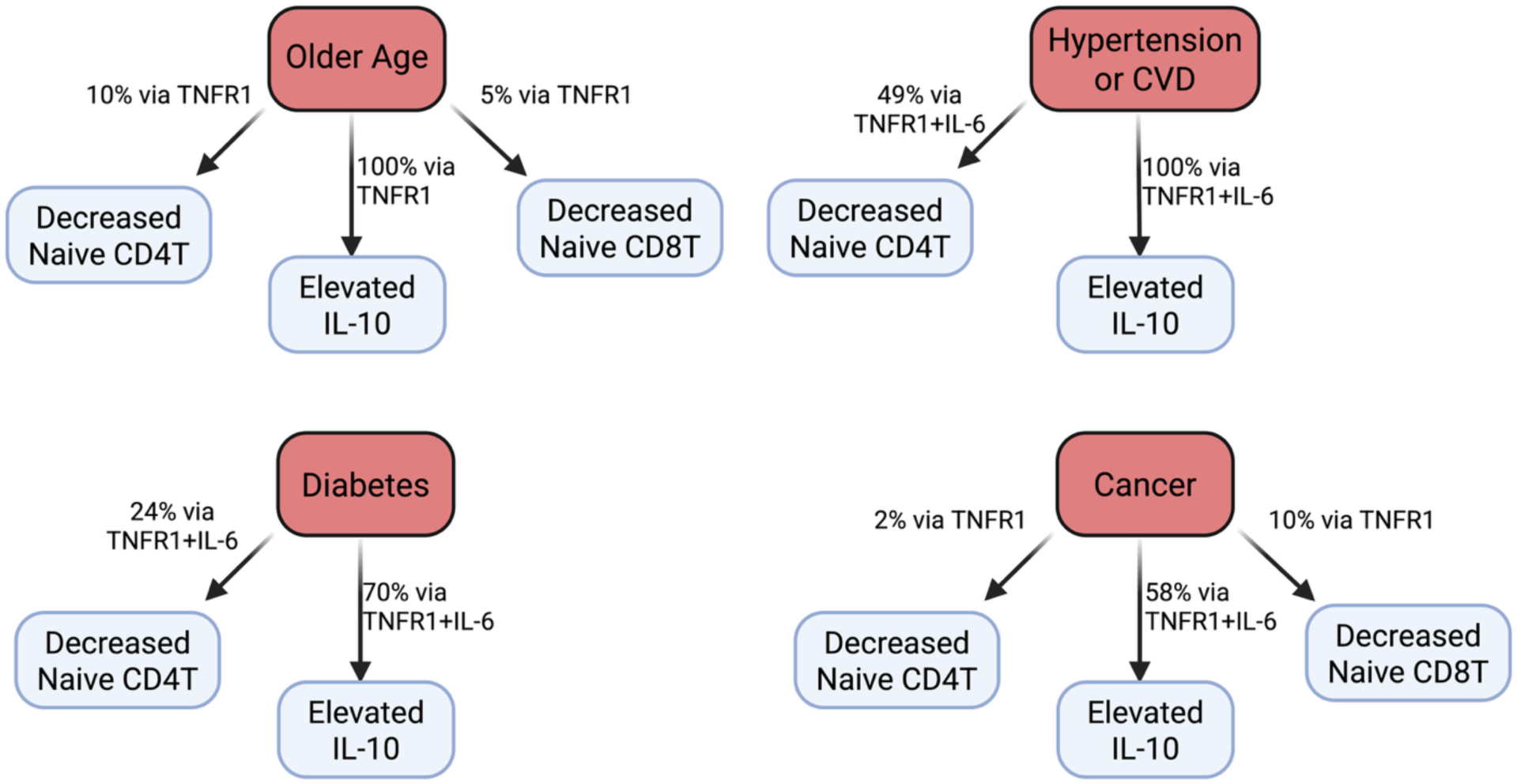
Key mediation results in the US HRS

Models predicting IL-10 and GDF15 in the UKB are shown in **Supplementary Table S7**. In the base model without pro-inflammatory cytokines, older age and chronic diseases were positively associated with IL-10 and GDF15. Consistent with mediation, TNFR1, IL-6, and TNF-α were all positively associated with IL-10 and GDF15, while their addition to the model attenuated age and disease associations. Associations between cytokines and odds of hospitalization or death from COVID-19 in the UKB are shown in **Table 3**. Older age, as well as diseases of hypertension or CVD and diabetes were associated with higher odds of hospitalization or death. Consistent with mediation, both IL-6 and TNFR1 were positively associated with odds of hospitalization or death, while their addition to the model attenuated the age and disease associations. Specifically, a standard deviation increase in TNFR1 was associated with 46% higher odds of hospitalization or death, while for IL-6 it was 28%. Also consistent with mediation, both IL-10 and GDF15 were positively associated with odds of hospitalization or death, and their addition to the model attenuated both age and disease associations as well as the associations with IL-6 and TNFR1. Specifically, a standard deviation increase in IL-10 was associated with 12% higher odds of hospitalization or death, while for GDF15 it was 45%. In the final model with all predictors, VIF’s were the following: age = 1.22, sex = 1.04, hypertension or CVD = 1.18, diabetes = 1.26, cancer = 1.01, IL-6 = 1.27, TNFR1 = 1.99, IL-1β = 1.04, TNF-α = 1.55, IL-10 = 1.11, GDF15 = 2.16.

**Table 3.** Logistic regression models predicting odds of hospitalization or death from COVID-19 (n = 552 out of 40,638). Cytokine coefficients reflect changes in standard deviations. Estimates are reported as Odds Ratios (OR) with 95% confidence intervals (LCI, UCI).

|  | OR with 95% confidence intervals |  |  |
| --- | --- | --- | --- |
| Age (years) | 1.05*** (1.04, 1.06) | 1.03*** (1.02, 1.04) | 1.02** (1.01, 1.03) |
| Male (0,1) | 1.24* (1.03, 1.45) | 1.21* (1.00, 1.41) | 1.14 (0.94, 1.33) |
| Hypertension or CVD (0,1) | 1.62*** (1.33, 1.91) | 1.33** (1.09, 1.58) | 1.28** (1.04, 1.51) |
| Diabetes (0,1) | 1.94*** (1.41, 2.46) | 1.50** (1.09, 1.92) | 1.06 (0.75, 1.38) |
| Cancer (0,1) | 1.10 (0.77, 1.43) | 1.02 (0.71, 1.33) | 1.00 (0.70, 1.30) |
| IL-6 |  | 1.28*** (1.17, 1.40) | 1.23*** (1.12, 1.35) |
| TNFR1 |  | 1.46*** (1.31, 1.61) | 1.26*** (1.12, 1.40) |
| IL-1β | | 0.91* (0.83, 0.99) | 0.92 (0.83, 1.00) |
| TNF-α | | 1.06 (0.94, 1.18) | 0.99 (0.87, 1.11) |
| IL-10 |  |  | 1.12* (1.02, 1.22) |
| GDF15 |  |  | 1.45*** (1.29, 1.61) |

\* p < 0.05; \*\* p < 0.01; \*\*\* p < 0.001

Since risk of hospitalization or death from COVID-19 could be due to a combination of an energy conservation response as well as dysfunctional immunity, we considered a sensitivity analysis excluding individuals with diseases or conditions related to dysregulated inflammation. We excluded individuals with hypertension or CVD, diabetes, and cancer, as well as those with COPD, asthma, or allergies. This resulted in a subsample of 17,665 in which 154 were later hospitalized or died from COVID-19. Results were largely similar compared to the full sample (**Supplementary Table S8**). TNFR1, IL-10, and GDF15 were still positively associated with odds of hospitalization or death, although IL-6 was not associated in the subsample.

**Figure 3.**
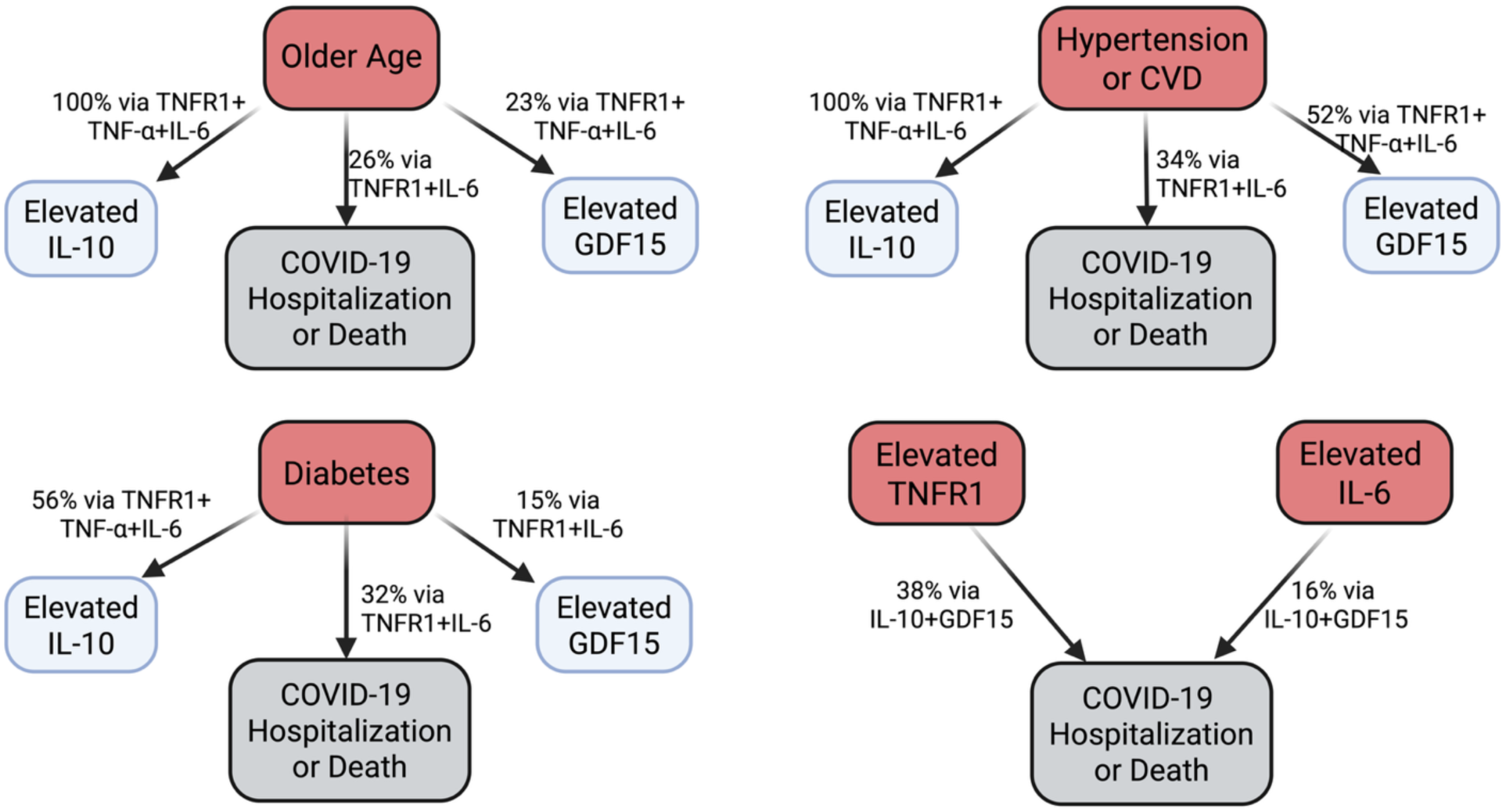
Key mediation results in the UKB

Mediation results for cytokines in the UKB are shown in **Supplementary Table S9**. Collectively, TNFR1, IL-6, and TNF-α mediated 100% of the positive association between age and IL-10, although none were significant independently. The positive association between age and GDF15 was mediated 11.0% by TNFR1, 2.2% by IL-6, and 1.2% by TNF-α independently. Similarly to age, TNFR1, IL-6, and TNF-α collectively mediated 100% of the positive association between hypertension or CVD and IL-10, although none were significant independently. The positive association between hypertension or CVD and GDF15 was mediated 27.9% by TNFR1, 13.6% by IL-6, and 1.9% by TNF-α independently. The positive association between diabetes and IL-10 was mediated 14.7% by TNFR1, 8.1% by IL-6, and 13.1% by TNF-α independently. The positive association between diabetes and GDF15 was mediated 8.0% by TNFR1 and 1.3% by IL-6 independently. The positive association between cancer and IL-10 was mediated 5.6% by TNFR1 and 4.1% by IL-6 independently, while the positive association for GDF15 was mediated 36.1% by TNFR1 and 10.5% by IL-6 independently.

Mediation results for cytokines and odds of hospitalization or death from COVID-19 in the UKB are shown in **Supplementary Table S10**. The positive association between age and risk of hospitalization or death was mediated 16.2% by TNFR1, 6.8% by IL-6, and 38.2% by GDF15. The positive association between hypertension or CVD and risk of hospitalization or death was mediated 16.4% by TNFR1 and 12.0% by IL-6, as well as 12.4% by GDF15. The positive association between diabetes and risk of hospitalization or death was mediated 26.7% by TNFR1, 8.3% by IL-6, and 81.6% by GDF15. Amongst the cytokines, the positive association between TNFR1 and risk of hospitalization or death was mediated 2.2% by IL-10 and 28.0% by GDF15. Finally, the positive association between IL-6 and risk of hospitalization or death was mediated 2.9% by IL-10 and 14.9% by GDF15. In the subsample without inflammatory diseases or conditions, the positive association between age and risk of hospitalization or death was mediated 12.1% by TNFR1 and 37.6% by GDF15 (**Supplementary Table S11**). The positive association between TNFR1 and risk of hospitalization or death was mediated 5.0% by IL-10 and 32.7% by GDF15.

## Discussion

Here we found evidence in two large human cohorts that is consistent with an energy conservation model of inflammaging and immunosenescence. TNFR1, an indicator of inflammaging, was inversely associated with naïve T cells in the HRS, positively associated with odds of hospitalization or death from COVID-19 in the UKB, and mediated the associations between older age and these outcomes (10% for CD4T, 5% for CD8T, 16% for COVID-19). We also found evidence consistent with the role of anti-inflammaging in augmenting immunosenescence in response to inflammaging. The anti-inflammatory cytokine IL-10 was positively associated with odds of hospitalization or death from COVID-19 and mediated 2% of the TNFR1 association. Finally, in further support of the energy conservation model of immune aging, GDF15, which is elevated in response to chronic inflammation and contributes to immune tolerance, also mediated 28% of the association between TNFR1 and increased risk of hospitalization or death, as well as 38% of the age-related increased risk independent of TNFR1.

Our findings are consistent with multiple previous studies on inflammation and immune function. For example, in the Multi-Ethnic Study of Atherosclerosis (MESA), both subclinical atherosclerosis and IL-6 were inversely associated with naïve CD4T cells (Olson et al., 2013). Another study of older individuals also found an inverse association between IL-6 and naïve T cells (Merellano-Navarro et al., 2021). Further, both subclinical atherosclerosis and elevated C-Reactive Protein (CRP), a marker of chronic low-grade inflammation, were found to be inversely associated with the inflammatory response to vaccination (McDade, Borja, Kuzawa, Perez, & Adair, 2015; McDade et al., 2020). Further, CRP has also been found to be inversely associated with the antibody response to vaccination (McDade, Adair, Feranil, & Kuzawa, 2011).

Adaptive remodeling in later life to promote short term survival raises the question of how this response would evolve with diminishing selection pressure. One potential explanation is that this energy conservation response evolved to function during acute periods of somatic damage in early life. An example of this is acute or transient thymus involution, which can be induced by a variety of exposures inducing somatic damage or energetic stress (e.g., chemotherapy, infection, malnutrition, pregnancy) (Ansari & Liu, 2017; Granadier, Acenas, & Dudakov, 2025). There is also evidence that several cytokines play a role in acute thymus involution, including the ones examined here (Ansari & Liu, 2017; Granadier et al., 2025). This acute response could have plausibly been co-opted for later life. Further, while selection pressure wanes, it is not completely absent, and this could further reinforce the energy conservation response to inflammaging.

A key open question is where compensation ends and pathology begins, i.e., how to empirically distinguish adaptive remodeling from maladaptive decline in late life (Fülöp et al., 2020). Our approach here was to assess how robust th associations were between age, cytokines, and COVID-19 health risk after excluding individuals with reported inflammatory diseases or conditions. While this approach is admittedly imperfect, if associations attenuated when individuals with the most dysregulated inflammatory responses were removed, it would suggest dysregulation was playing a larger role. This appeared to be the case for IL-6, which was no longer predictive of COVID-19 health after removing individuals with inflammatory conditions. In the HRS, IL-6 was more strongly implicated disease-related decreases in naïve T cells versus age-related, further suggesting dysregulated inflammation is playing a larger role in the IL-6 associations. However, associations for TNFR1, GDF15, and IL-10 in the UKB were either very similar or slightly strengthened after removing individuals with inflammatory conditions, suggesting dysregulated inflammation is not heavily confounding these relationships.

While here we propose that the BEC model can help explain atrophy of the immune repertoire, it is likely not a comprehensive explanation. With decreasing residual life expectancy, the benefits of investment in the immune repertoire decline. As a result, we should expect atrophy of the immune repertoire with declining life expectancy, as is the case in later life. This might explain why we found strong inverse associations between male sex, which is associated with lower life expectancy, and naïve T cells. Relatedly, most infectious exposures have already been encountered by later life, and as a result the marginal value of further investment in the immune repertoire declines.

Our results for IL-10 are consistent with prior studies suggesting this cytokine plays a role in augmenting immunosenescence and increased vulnerability to infection in later life (Almanan et al., 2020; McElhaney et al., 2012). However, its influence on disease and aging is complex and context specific. Current evidence suggests that greater IL-10 is protective from the development of inflammaging and potentially some inflammatory diseases such as CVD (Dagdeviren et al., 2016; Li et al., 2024; McElhaney et al., 2012; Minciullo et al., 2016). Further, the increase in IL-10 in response to inflammaging might serve multiple functions. In addition to energy conservation, IL-10 might also play a role in striking a balance between an excessive inflammatory response that contributes to aging and disease and an impaired immune response to infections (Hedrich & Bream, 2010; Huang et al., 2020; Ouyang, Rutz, Crellin, Valdez, & Hymowitz, 2011). GDF15 has also been proposed to serve this function (Moon et al., 2020)

While here we use the terminology of inflammaging and anti-inflammaging, consistent with pro- and anti-inflammatory, it is important to note these cytokines likely reflect a single immune response. Evidence for this can be seen for example in the positive correlations amongst cytokines with opposing functions reported here and in previous studies, as well as similar principal component loadings (Aronoff et al., 2025; Franck, Tanner, et al., 2025; Morrisette-Thomas et al., 2014).

Our findings also highlight TNFR1 as an important marker of inflammaging. This is consistent with a recent study using a battery of cytokines to measure inflammaging across human populations. At least among industrialized populations, TNFR1 was a strong indicator of inflammaging (Franck, Tanner, et al., 2025). In contrast, we found that other markers often used to measure inflammaging, including IL-6, IL-1β, and TNF-α, showed weaker associations with naïve T cells or odds of hospitalization or death from COVID-19. These results highlight the importance of measuring TNFR1 as an indicator of inflammaging in future studies.

This energy conservation model linking inflammaging to immunosenescence has important health implications. For example, it can offer an explanation of the rapamycin immune function “paradox”. While rapamycin was originally used as an immunosuppressant for organ transplant patients, it has been found to improve antibody responses to vaccination as well as reduce infection risk over follow-up (Mannick et al., 2014; Mannick et al., 2018). The energy conservation model of immune aging suggests this effect is likely indirect. Rapamycin can inhibit SASP signaling, thereby reducing inflammaging (Blagosklonny, 2022; Sorrenti et al., 2022; R. Wang, Sunchu, & Perez, 2017). As a result, the divestment from the immune repertoire could be reversed, improving immune responses to novel infectious exposures (Aronoff & Trumble, 2025). Relatedly, previous findings of transient and reversible thymus involution highlighted above further suggests the potential for reversing immunosenescence if inflammaging can be attenuated. This could be accomplished therapeutically, such as with rapamycin, or through lifestyle changes. For example, among older adults, physical activity has been found to be inversely associated with IL-6 and positively associated with naïve CD4T cells (Merellano-Navarro et al., 2021). Further, non-industrialized human populations that are highly physically active with minimal excess caloric consumption, such as Tsimane forager-horticulturalists in the Bolivian Amazon, and the Orang Asli show markedly attenuated inflammaging (Aronoff et al., 2025; Franck, Tanner, et al., 2025). Finally, a horticulture lifestyle intervention showed a reduction in IL-6 and increase in naïve CD8T cells (Wong et al., 2021). However, it is important to distinguish between interventions that might artificially dampen inflammaging such as anti-inflammatory medications and interventions that might address the causes of inflammaging, such as physical activity. Under the BEC, the former might succeed in reducing immunosenescence only at the cost of perturbing the complex energetic balance the body is attempting to maintain during aging.

Our results also shed light on the relationship between comorbidities and health risk from COVID-19 (Bucholc et al., 2022). While a dysregulated inflammatory response is likely an important link between chronic disease and COVID-19 health risk, another potential pathway is energy conservation-driven immunosenescence. These processes could work synergistically. Viruses can enhance their proliferation in the host due to impaired immunosurveillance, and once this heavy viral load is detected it elicits an excessive inflammatory response that induces host damage (e.g., “cytokine storm”) (Yang et al., 2021).

Our study is not without limitations. While we assessed indicators of both investment in the immune repertoire and an outcome of vulnerability to novel infection, these were measured in different individuals across the two US and UK cohorts. Measuring these outcomes in the same individuals, as well as assessing them longitudinally, is needed. In addition, our measures of cytokines were cross-sectional and part of an observational study design. As a result, while the mediation analysis was motivated by and consistent with previous experimental studies, we cannot definitively assess causal ordering. Further, due to the complex signaling network of cytokines that results in a versatile emergent property of immune function (Franck, Daunizeau, et al., 2025), we also cannot rule out bidirectional causality amongst these cytokines.

In conclusion, here we find evidence consistent with an energy conservation model of immune aging. Immunosenescence, at least in part, can be understood as an energy conservation response to rising inflammaging in later life. This response likely promotes short term survival at the expense of greater vulnerability to novel infections in the future. Considering waning selection pressure with age, this response likely originally evolved for surviving short term stressors in early life and has been co-opted to operate chronically in later life. Finally, our study also highlights the role of anti-inflammatory signaling, or anti-inflammaging, as a contributor to immunosenescence.

## Acknowledgements

This research has been conducted using the UK Biobank Resource under the approved application number 772113. We are grateful to the US HRS and UKB study participants, as well as the researchers who have created and maintained these publicly available databases. Partial funding support came from the NIH/National Institute on Aging (R01AG054442). Figures were created in Biorender.

## Author contributions

Jacob E. Aronoff: conceptualization (lead), statistical analysis, writing (original draft). Maximilien Franck: conceptualization (supporting), writing (review and editing). Alan A. Cohen: conceptualization (supporting), writing (review and editing). Benjamin C. Trumble: conceptualization (supporting), writing (review and editing).

## References

Ahmed, D. S., Isnard, S., Berini, C., Lin, J., Routy, J.-P., & Royston, L. (2022). Coping with stress: the mitokine GDF-15 as a biomarker of COVID-19 severity. Frontiers in immunology, 13, 820350.

Almanan, M., Raynor, J., Ogunsulire, I., Malyshkina, A., Mukherjee, S., Hummel, S. A., . . . Alenghat, T. (2020). IL-10–producing Tfh cells accumulate with age and link inflammation with age-related immune suppression. Science advances, 6(31), eabb0806.

Ansari, A. R., & Liu, H. (2017). Acute thymic involution and mechanisms for recovery. Archivum Immunologiae et Therapiae Experimentalis, 65(5), 401–420.

Aronoff, J. E., Jenkins, C. L., Garcia, A. R., Koebele, S. V., Ghafoor, S., Woolard, K. L., . . . Beheim, B. (2025). Inflammaging is minimal among forager-horticulturalists in the Bolivian Amazon. Proceedings B, 292(2053), 20251111.

Aronoff, J. E., & Trumble, B. C. (2025). An evolutionary medicine and life history perspective on aging and disease: Trade-offs, hyperfunction, and mismatch. *Evolution*, Medicine, and Public Health, 13(1), 111–124.

Blackwell, A. D., Trumble, B. C., Maldonado Suarez, I., Stieglitz, J., Beheim, B., Snodgrass, J. J., . . . Gurven, M. (2016). Immune function in Amazonian horticulturalists. Annals of Human Biology, 43(4), 382–396.

Blagosklonny, M. V. (2022). Cell senescence, rapamycin and hyperfunction theory of aging. Cell Cycle, 21(14), 1456–1467.

Bu, S., Royston, L., Mabanga, T., Berini, C. A., Tremblay, C., Lebouché, B., . . . Isnard, S. (2024). Proteomics validate circulating GDF-15 as an independent biomarker for COVID-19 severity. Frontiers in immunology, 15, 1377126.

Bucholc, M., Bradley, D., Bennett, D., Patterson, L., Spiers, R., Gibson, D., . . . Bjourson, A. J. (2022). Identifying pre-existing conditions and multimorbidity patterns associated with in-hospital mortality in patients with COVID-19. Scientific reports, 12(1), 17313.

Clancy, K. B., Klein, L. D., Ziomkiewicz, A., Nenko, I., Jasienska, G., & Bribiescas, R. G. (2013). Relationships between biomarkers of inflammation, ovarian steroids, and age at menarche in a rural Polish sample. American Journal of Human Biology, 25(3), 389–398.

Coppé, J.-P., Patil, C. K., Rodier, F., Sun, Y., Muñoz, D. P., Goldstein, J., . . . Campisi, J. (2008). Senescence-associated secretory phenotypes reveal cell-nonautonomous functions of oncogenic RAS and the p53 tumor suppressor. PLoS biology, 6(12), e301.

Crimmins, E., Faul, J., Thyagarajan, B., & Weir, D. (2017). Venous blood collection and assay protocol in the 2016 Health and Retirement Study 2016 Venous Blood Study (VBS). Retrieved from

Dagdeviren, S., Jung, D. Y., Friedline, R. H., Noh, H. L., Kim, J. H., Patel, P. R., . . . Hu, X. (2016). IL-10 prevents aging-associated inflammation and insulin resistance in skeletal muscle. The FASEB Journal, 31(2), 701.

Franceschi, C., Bonafè, M., Valensin, S., Olivieri, F., De Luca, M., Ottaviani, E., & De Benedictis, G. (2000). Inflamm-aging: an evolutionary perspective on immunosenescence. Annals of the new York Academy of Sciences, 908(1), 244–254.

Franceschi, C., Capri, M., Monti, D., Giunta, S., Olivieri, F., Sevini, F., . . . Scurti, M. (2007). Inflammaging and anti-inflammaging: a systemic perspective on aging and longevity emerged from studies in humans. Mechanisms of ageing and development, 128(1), 92–105.

Franck, M., Daunizeau, C., Aronoff, J. E., Tanner, K., Trumble, B. C., Franceschi, C., . . . Gurven, M. (2025). Inflamm-aging as a diverse and context-dependent process: from species and population differences to individual trajectories. Ageing Research Reviews, 102880.

Franck, M., Tanner, K. T., Tennyson, R. L., Daunizeau, C., Ferrucci, L., Bandinelli, S., . . . Stieglitz, J. (2025). Nonuniversality of inflammaging across human populations. Nature Aging, 1–10.

Fülöp, T., Larbi, A., Dupuis, G., Le Page, A., Frost, E. H., Cohen, A. A., . . . Franceschi, C. (2018). Immunosenescence and inflamm-aging as two sides of the same coin: friends or foes? Frontiers in immunology, 8, 1960.

Fülöp, T., Larbi, A., Hirokawa, K., Cohen, A., & Witkowski, J. (2020). Immunosenescence is both functional/adaptive and dysfunctional/maladaptive. Paper presented at the Seminars in Immunopathology.

Fülöp, T., Larbi, A., Pawelec, G., Khalil, A., Cohen, A., Hirokawa, K., . . . Franceschi, C. (2021). Immunology of aging: the birth of inflammaging. Clinical reviews in allergy & immunology, 1–14.

Granadier, D., Acenas, D., & Dudakov, J. A. (2025). Endogenous thymic regeneration: restoring T cell production following injury. Nature reviews immunology, 1–18.

Hedrich, C. M., & Bream, J. H. (2010). Cell type-specific regulation of IL-10 expression in inflammation and disease. Immunologic research, 47(1), 185–206.

Huang, Y.-H., Chen, M.-H., Guo, Q.-L., Chen, Z.-X., Chen, Q.-D., & Wang, X.-Z. (2020). Interleukin-10 induces senescence of activated hepatic stellate cells via STAT3-p53 pathway to attenuate liver fibrosis. Cellular signalling, 66, 109445.

Hunter-Schlichting, D., Lane, J., Cole, B., Flaten, Z., Barcelo, H., Ramasubramanian, R., . . . Pankratz, N. (2020). Validation of a hybrid approach to standardize immunophenotyping analysis in large population studies: The Health and Retirement Study. Scientific reports, 10(1), 8759.

Imai, K., Keele, L., & Tingley, D. (2010). A general approach to causal mediation analysis. Psychological methods, 15(4), 309.

Ip, W. E., Hoshi, N., Shouval, D. S., Snapper, S., & Medzhitov, R. (2017). Anti-inflammatory effect of IL-10 mediated by metabolic reprogramming of macrophages. Science, 356(6337), 513–519.

Li, Y., Yin, H., Yuan, H., Wang, E., Wang, C., Li, H., . . . Bai, J. (2024). IL-10 deficiency aggravates cell senescence and accelerates BLM-induced pulmonary fibrosis in aged mice via PTEN/AKT/ERK pathway. BMC Pulmonary Medicine, 24(1), 443.

Lochmiller, R. L., & Deerenberg, C. (2000). Trade-offs in evolutionary immunology: just what is the cost of immunity? Oikos, 88(1), 87–98.

Luan, H. H., Wang, A., Hilliard, B. K., Carvalho, F., Rosen, C. E., Ahasic, A. M., . . . Yu, S. (2019). GDF15 is an inflammation-induced central mediator of tissue tolerance. Cell, 178(5), 1231–1244. e1211.

Mannick, J. B., Del Giudice, G., Lattanzi, M., Valiante, N. M., Praestgaard, J., Huang, B., . . . Carson, S. (2014). mTOR inhibition improves immune function in the elderly. Science translational medicine, 6(268), 268ra179–268ra179.

Mannick, J. B., Morris, M., Hockey, H.-U. P., Roma, G., Beibel, M., Kulmatycki, K., . . . Quinn, D. (2018). TORC1 inhibition enhances immune function and reduces infections in the elderly. Science translational medicine, 10(449), eaaq1564.

McDade, T. W. (2003). Life history theory and the immune system: steps toward a human ecological immunology. American Journal of Physical Anthropology: The Official Publication of the American Association of Physical Anthropologists, 122(S37), 100–125.

McDade, T. W., Adair, L., Feranil, A. B., & Kuzawa, C. (2011). Positive antibody response to vaccination in adolescence predicts lower C-reactive protein concentration in young adulthood in the Philippines. American Journal of Human Biology, 23(3), 313–318.

McDade, T. W., Borja, J. B., Kuzawa, C. W., Perez, T. L. L., & Adair, L. S. (2015). C-reactive protein response to influenza vaccination as a model of mild inflammatory stimulation in the Philippines. Vaccine, 33(17), 2004–2008.

McDade, T. W., Borja, J. B., Lee, N., Aquino, C. T., Barrett, T., Adair, L. S., & Kuzawa, C. W. (2020). C-reactive protein response to influenza vaccination predicts cardiovascular disease risk in the Philippines. Biodemography and Social Biology, 65(1), 88–96.

McDade, T. W., Georgiev, A. V., & Kuzawa, C. W. (2016). Trade-offs between acquired and innate immune defenses in humans. Evolution, Medicine, and Public Health, 2016(1), 1–16.

McElhaney, J. E., Zhou, X., Talbot, H. K., Soethout, E., Bleackley, R. C., Granville, D. J., & Pawelec, G. (2012). The unmet need in the elderly: how immunosenescence, CMV infection, co-morbidities and frailty are a challenge for the development of more effective influenza vaccines. Vaccine, 30(12), 2060–2067.

Merellano-Navarro, E., Olate-Briones, A., Norambuena-Mardones, L., Rojas-Ramos, V., de la Plata-Luna, A. M., Faúndez-Acuña, J. Y., . . . Herrada, A. A. (2021). Reduced Naïve T Cell Numbers Correlate with Increased Low-Grade Systemic Inflammation During Ageing and Can be Modulated by Physical Activity. International Journal of Morphology, 39(3).

Minciullo, P. L., Catalano, A., Mandraffino, G., Casciaro, M., Crucitti, A., Maltese, G., . . . Basile, G. (2016). Inflammaging and anti-inflammaging: the role of cytokines in extreme longevity. Archivum Immunologiae et Therapiae Experimentalis, 64, 111–126.

Moon, J. S., Goeminne, L. J., Kim, J. T., Tian, J. W., Kim, S. H., Nga, H. T., . . . Lee, Y. S. (2020). Growth differentiation factor 15 protects against the aging-mediated systemic inflammatory response in humans and mice. Aging Cell, 19(8), e13195.

Morrisette-Thomas, V., Cohen, A. A., Fülöp, T., Riesco, É., Legault, V., Li, Q., . . . Ferrucci, L. (2014). Inflamm-aging does not simply reflect increases in pro-inflammatory markers. Mechanisms of ageing and development, 139, 49–57.

Muehlenbein, M. P., Hirschtick, J. L., Bonner, J. Z., & Swartz, A. M. (2010). Toward quantifying the usage costs of human immunity: altered metabolic rates and hormone levels during acute immune activation in men. American Journal of Human Biology: The Official Journal of the Human Biology Association, 22(4), 546–556.

Olson, N. C., Doyle, M. F., Jenny, N. S., Huber, S. A., Psaty, B. M., Kronmal, R. A., & Tracy, R. P. (2013). Decreased naive and increased memory CD4+ T cells are associated with subclinical atherosclerosis: the multi-ethnic study of atherosclerosis. PloS one, 8(8), e71498.

Ouyang, W., Rutz, S., Crellin, N. K., Valdez, P. A., & Hymowitz, S. G. (2011). Regulation and functions of the IL-10 family of cytokines in inflammation and disease. Annual review of immunology, 29(1), 71–109.

Partridge, L., Fuentealba, M., & Kennedy, B. K. (2020). The quest to slow ageing through drug discovery. Nature Reviews Drug Discovery, 19(8), 513–532.

Paul, W. (2013). Fundamental Immunology. Philadelphia. In: Lippincott Williams & Wilkins, a Wolters Kluwer business.

Pence, B. D. (2022). Growth differentiation factor-15 in immunity and aging. Frontiers in aging, 3, 837575.

Peters, A., Delhey, K., Nakagawa, S., Aulsebrook, A., & Verhulst, S. (2019). Immunosenescence in wild animals: meta-analysis and outlook. Ecology Letters, 22(10), 1709–1722.

Pontzer, H., Yamada, Y., Sagayama, H., Ainslie, P. N., Andersen, L. F., Anderson, L. J., . . . Blaak, E. E. (2021). Daily energy expenditure through the human life course. Science, 373(6556), 808–812.

Roff, D. A. (2002). Life history evolution.

Salminen, A. (2025). GDF15/MIC-1: A stress-induced immunosuppressive factor which promotes the aging process. Biogerontology, 26(1), 19.

Shanley, D. P., Aw, D., Manley, N. R., & Palmer, D. B. (2009). An evolutionary perspective on the mechanisms of immunosenescence. Trends in immunology, 30(7), 374–381.

Shaulson, E. D., Cohen, A. A., & Picard, M. (2024). The brain–body energy conservation model of aging. Nature Aging, 1–18.

Sorrenti, V., Benedetti, F., Buriani, A., Fortinguerra, S., Caudullo, G., Davinelli, S., . . . Scapagnini, G. (2022). Immunomodulatory and antiaging mechanisms of resveratrol, rapamycin, and metformin: focus on mTOR and AMPK signaling networks. Pharmaceuticals, 15(8), 912.

Stearns, S. C. (1992). The evolution of life histories.

Sun, B. B., Chiou, J., Traylor, M., Benner, C., Hsu, Y.-H., Richardson, T. G., . . . Vasquez-Grinnell, S. G. (2023). Plasma proteomic associations with genetics and health in the UK Biobank. Nature, 622(7982), 329–338.

Thyagarajan, B., Barcelo, H., Crimmins, E., Weir, D., Minnerath, S., Vivek, S., & Faul, J. (2018). Effect of delayed cell processing and cryopreservation on immunophenotyping in multicenter population studies. Journal of immunological methods, 463, 61–70.

Thyagarajan, B., Faul, J., Vivek, S., Kim, J. K., Nikolich-Žugich, J., Weir, D., & Crimmins, E. M. (2022). Age-related differences in T-cell subsets in a nationally representative sample of people older than age 55: findings from the health and retirement study. The Journals of Gerontology: Series A, 77(5), 927–933.

Tingley, D., Yamamoto, T., Hirose, K., Keele, L., & Imai, K. (2014). Mediation: R package for causal mediation analysis.

Urlacher, S. S., Ellison, P. T., Sugiyama, L. S., Pontzer, H., Eick, G., Liebert, M. A., . . . Snodgrass, J. J. (2018). Tradeoffs between immune function and childhood growth among Amazonian forager-horticulturalists. Proceedings of the National Academy of Sciences, 115(17), E3914–E3921.

Wang, A., Luan, H. H., & Medzhitov, R. (2019). An evolutionary perspective on immunometabolism. Science, 363(6423), eaar3932.

Wang, R., Sunchu, B., & Perez, V. I. (2017). Rapamycin and the inhibition of the secretory phenotype. Experimental gerontology, 94, 89–92.

Wong, G. C. L., Ng, T. K. S., Lee, J. L., Lim, P. Y., Chua, S. K. J., Tan, C., . . . Sia, A. (2021). Horticultural therapy reduces biomarkers of immunosenescence and inflammaging in community-dwelling older adults: a feasibility pilot randomized controlled trial. The Journals of Gerontology: Series A, 76(2), 307–317.

Yang, L., Xie, X., Tu, Z., Fu, J., Xu, D., & Zhou, Y. (2021). The signal pathways and treatment of cytokine storm in COVID-19. Signal transduction and targeted therapy, 6(1), 255.

Zuk, M., & Stoehr, A. M. (2002). Immune defense and host life history. the american naturalist, 160(S4), S9–S22.

