## Supplemental Tables for "Evidence for an energetic trade-off model linking inflammaging and immunosenescence in the US Health and Retirement Study and UK Biobank"

**Supplementary Materials**

**Table S1.** Correlations among cytokines in the US HRS (n = 8,261)

| | IL-6 | TNFR1 |
|---|---|---|
| TNFR1 | 0.42 | |
| IL-10 | 0.35 | 0.44 |

All $p < 0.001$

**Table S2.** Correlations among cytokines in the UKB (n = 40,638)

| | IL-6 | TNFR1 | IL-10 | IL-1β | TNF-α |
|---|---|---|---|---|---|
| TNFR1 | 0.40 | | | | |
| IL-10 | 0.12 | 0.15 | | | |
| IL-1β | 0.08 | 0.14 | 0.04 | | |
| TNF-α | 0.29 | 0.45 | 0.24 | 0.16 | |
| GDF15 | 0.35 | 0.53 | 0.08 | 0.06 | 0.34 |

All $p < 0.001$

**Table S3.** HRS pro-inflammatory cytokine associations (n = 8,261)

| | IL-6 | TNFR1 |
|---|---|---|
| Age (years) | 0.015*** (0.001) | 0.033*** (0.001) |
| Male (0,1) | 0.038 (0.020) | -0.020 (0.018) |
| Hypertension or CVD (0,1) | 0.255*** (0.022) | 0.293*** (0.019) |
| Diabetes (0,1) | 0.233*** (0.024) | 0.322*** (0.021) |
| Cancer (0,1) | 0.094*** (0.027) | 0.130*** (0.024) |

*** $p < 0.001$

**Table S4.** UKB pro-inflammatory cytokine associations (n = 40,638)

| | IL-6 | TNFR1 | IL-1β | TNF-α |
|---|---|---|---|---|
| Age (years) | 0.018*** (0.001) | 0.026*** (0.001) | 0.001* (0.001) | 0.017*** (0.001) |
| Male (0,1) | -0.018* (0.009) | 0.129*** (0.009) | -0.063*** (0.009) | -0.053*** (0.008) |
| Hypertension or CVD (0,1) | 0.264*** (0.010) | 0.262*** (0.010) | 0.072*** (0.011) | 0.151*** (0.009) |
| Diabetes (0,1) | 0.253*** (0.021) | 0.387*** (0.021) | 0.010 (0.022) | 0.197*** (0.019) |
| Cancer (0,1) | 0.076*** (0.017) | 0.099*** (0.017) | -0.0002 (0.018) | 0.037* (0.016) |

* $p < 0.05$; ** $p < 0.01$; *** $p < 0.001$

**Table S5.** Regression models predicting IL-10 and Naïve T cells in the US HRS, adjusting for CMV antibodies. Cytokines, T cell counts, and CMV antibodies were standardized, with resulting regression coefficients reflecting changes in standard deviations (n = 8,261). Estimates are reported as β (SE).

| | IL-10 | IL-10 | CD4T | CD4T | CD8T | CD8T |
|---|---|---|---|---|---|---|
| Age (years) | -0.003* (0.001) | -0.003* (0.001) | -0.015*** (0.001) | -0.013*** (0.001) | -0.034*** (0.001) | -0.034*** (0.001) |
| Male (0,1) | 0.098*** (0.018) | 0.097*** (0.018) | -0.358*** (0.021) | -0.390*** (0.021) | -0.310*** (0.017) | -0.302*** (0.017) |
| Hypertension or CVD (0,1) | 0.026 (0.020) | 0.027 (0.020) | -0.027 (0.024) | -0.019 (0.023) | 0.003 (0.020) | 0.001 (0.020) |
| Diabetes (0,1) | 0.070** (0.022) | 0.071*** (0.022) | -0.087*** (0.025) | -0.069** (0.025) | 0.011 (0.021) | 0.007 (0.021) |
| Cancer (0,1) | 0.047 (0.024) | 0.046 (0.024) | -0.259*** (0.029) | -0.273*** (0.029) | -0.059* (0.024) | -0.055* (0.024) |
| IL-6 | 0.184*** (0.011) | 0.184*** (0.011) | -0.034** (0.013) | -0.028* (0.013) | 0.006 (0.011) | 0.004 (0.011) |
| TNFR1 | 0.366*** (0.012) | 0.366*** (0.012) | -0.061*** (0.015) | -0.062*** (0.014) | -0.064*** (0.012) | -0.064*** (0.012) |
| CMV Ab | | -0.005 (0.009) | | -0.130*** (0.010) | | 0.033*** (0.008) |

* $p < 0.05$; ** $p < 0.01$; *** $p < 0.001$

**Table S6.** Mediation results for age, disease (HCVD = hypertension or CVD), cytokines, and naïve T cells in the US HRS (n = 8,261)

| Mediation path | Prop. mediated | 95% CI | p |
|---|---|---|---|
| Age→TNFR1→IL-10 | 100 | 100 – 100 | <0.001 |
| HCVD→IL-6→IL-10 | 47.9 | 24.5 – 100 | 0.014 |
| HCVD→TNFR1→IL-10 | 74.2 | 52.5 – 100 | <0.001 |
| Diabetes→IL-6→IL-10 | 20.8 | 11.0 – 40.0 | <0.001 |
| Diabetes→TNFR1→IL-10 | 56.5 | 43.9 – 76.7 | <0.001 |
| Cancer→IL-6→IL-10 | 13.2 | 0.0 – 62.6 | 0.130 |
| Cancer→TNFR1→IL-10 | 43.2 | 22.5 – 97.5 | <0.001 |
| Age→TNFR1→CD4T | 10.1 | 5.7 – 15.0 | <0.001 |
| HCVD→IL-6→CD4T | 12.1 | 0.0 – 100 | 0.180 |
| HCVD→TNFR1→CD4T | 30.7 | 0.0 – 100 | 0.084 |
| Diabetes→IL-6→CD4T | 3.6 | 0.7 – 10.0 | 0.014 |
| Diabetes→TNFR1→CD4T | 14.9 | 6.7 – 32.0 | <0.001 |
| Cancer→TNFR1→CD4T | 2.3 | 1.0 – 4.0 | <0.001 |
| Age→TNFR1→CD8T | 5.0 | 3.2 – 7.0 | <0.001 |
| Cancer→TNFR1→CD8T | 9.6 | 3.7 – 31.0 | 0.002 |

**Table S7.** Regression models predicting IL-10 and GDF15 in the UKB. Cytokines were standardized, with resulting regression coefficients reflecting changes in standard deviations (n = 40,638). Estimates are reported as β (SE).

| | IL-10 | IL-10 | GDF15 | GDF15 |
|---|---|---|---|---|
| Age (years) | $0.002^{**}$ (0.001) | $-0.004^{***}$ (0.001) | $0.050^{***}$ (0.0005) | $0.038^{***}$ (0.0004) |
| Male (0,1) | $-0.025^{**}$ (0.009) | $-0.017^{*}$ (0.009) | $0.160^{***}$ (0.007) | $0.125^{***}$ (0.006) |
| Hypertension or CVD (0,1) | $0.032^{**}$ (0.010) | $-0.026^{**}$ (0.010) | $0.228^{***}$ (0.008) | $0.109^{***}$ (0.007) |
| Diabetes (0,1) | $0.134^{***}$ (0.021) | $0.059^{**}$ (0.021) | $1.050^{***}$ (0.017) | $0.888^{***}$ (0.015) |
| Cancer (0,1) | $0.063^{***}$ (0.018) | $0.046^{**}$ (0.017) | $0.078^{***}$ (0.014) | $0.036^{**}$ (0.012) |
| IL-6 | | $0.047^{***}$ (0.005) | | $0.104^{***}$ (0.004) |
| TNFR1 | | $0.042^{***}$ (0.006) | | $0.309^{***}$ (0.004) |
| IL-1β | | -0.006 (0.005) | | $-0.015^{***}$ (0.003) |
| TNF-α | | $0.237^{***}$ (0.006) | | $0.081^{***}$ (0.004) |

**Table S8.** Logistic regression models predicting odds of hospitalization or death from COVID-19 in subsample without inflammatory diseases/conditions (n = 154 out of n = 17,665).

| | Odds ratios with 95% confidence intervals | | |
|---|---|---|---|
| Age (years) | $1.06^{***}$ (1.03, 1.08) | $1.04^{***}$ (1.02, 1.06) | $1.03^{*}$ (1.00, 1.05) |
| Male (0,1) | $1.84^{***}$ (1.25, 2.44) | $1.76^{***}$ (1.18, 2.33) | $1.64^{**}$ (1.10, 2.18) |
| IL-6 | | 1.14 (0.94, 1.35) | 1.06 (0.86, 1.26) |
| TNFR1 | | $1.45^{***}$ (1.15, 1.76) | 1.24 (0.96, 1.52) |
| IL-1β | | 0.91 (0.75, 1.07) | 0.92 (0.76, 1.09) |
| TNF-α | | 1.10 (0.87, 1.34) | 0.99 (0.77, 1.21) |
| IL-10 | | | $1.27^{**}$ (1.07, 1.48) |
| GDF15 | | | $1.53^{***}$ (1.19, 1.88) |

Listed diseases or conditions include hypertension or CVD, diabetes, cancer, COPD, asthma, or allergy. Cytokines were standardized, with resulting regression coefficients reflecting changes in standard deviations.

$^{*}$ $p < 0.05$; $^{**}$ $p < 0.01$; $^{***}$ $p < 0.001$

**Table S9.** Mediation results for age, diseases (HCVD = hypertension or CVD) and cytokines (n = 40,638)

| Mediation path | Proportion mediated | 95% CI | p |
|---|---|---|---|
| Age→IL-6→GDF15 | 2.2 | 1.8 – 2.5 | <0.001 |
| Age→TNFR1→GDF15 | 11.0 | 10.3 – 11.8 | <0.001 |
| Age→TNF-α→GDF15 | 1.2 | 1.0 – 1.5 | <0.001 |
| HCVD→IL-6→GDF15 | 13.6 | 11.5 – 16.0 | <0.001 |
| HCVD→TNFR1→GDF15 | 27.9 | 24.1 – 32.1 | <0.001 |
| HCVD→TNF-α→GDF15 | 1.9 | 0.7 – 3.2 | 0.002 |
| Diabetes→IL-6→IL-10 | 8.1 | 4.2 – 20.9 | <0.001 |
| Diabetes→TNFR1→IL-10 | 14.7 | 7.9 – 36.5 | <0.001 |
| Diabetes→TNF-α→IL-10 | 13.1 | 1.7 – 35.0 | 0.034 |
| Diabetes→IL-6→GDF15 | 1.3 | 0.9 – 1.7 | <0.001 |
| Diabetes→TNFR1→GDF15 | 8.0 | 7.0 – 9.2 | <0.001 |
| Cancer→IL-6→IL-10 | 4.1 | 1.0 – 15.1 | 0.012 |
| Cancer→TNFR1→IL-10 | 5.6 | 2.5 – 16.1 | 0.006 |
| Cancer→IL-6→GDF15 | 10.5 | 2.4 – 26.3 | 0.014 |
| Cancer→TNFR1→GDF15 | 36.1 | 21.3 – 63.0 | <0.001 |

**Table S10.** Mediation results for age, diseases (HCVD = hypertension or CVD), cytokines, and COVID-19 hospitalizations or deaths in the UKB (n = 40,638 with n = 552 hospitalized or died)

| Mediation path | Proportion mediated | 95% CI | p |
|---|---|---|---|
| Age→IL-6→COVID | 6.8 | 3.6 – 13.0 | <0.001 |
| Age→TNFR1→COVID | 16.2 | 10.1 – 27.7 | <0.001 |
| Age→GDF15→COVID | 38.2 | 21.3 – 77.8 | <0.001 |
| HCVD→IL-6→COVID | 12.0 | 6.0 – 33.8 | <0.001 |
| HCVD→TNFR1→COVID | 16.4 | 8.6 – 37.9 | <0.001 |
| HCVD→GDF15→COVID | 12.4 | 5.5 – 36.1 | 0.008 |
| Diabetes→IL-6→COVID | 8.3 | 2.7 – 53.0 | 0.026 |
| Diabetes→TNFR1→COVID | 26.7 | 13.3 – 1.00 | 0.014 |
| Diabetes→GDF15→COVID | 81.6 | 26.2 – 1.00 | 0.046 |
| TNFR1→IL-10→COVID | 2.2 | 0.5 – 5.9 | 0.008 |
| TNFR1→GDF15→COVID | 28.0 | 15.3 – 48.5 | <0.001 |
| IL-6→IL-10→COVID | 2.9 | 0.6 – 7.3 | 0.008 |
| IL-6→GDF15→COVID | 14.9 | 7.5 – 28.1 | <0.001 |

**Table S11.** Mediation results for age, cytokines, and COVID-19 hospitalizations or deaths in the UKB subsample without listed diseases (n = 17,665 total with n = 154 hospitalized or died)

| Mediation path | Proportion mediated | 95% CI | p |
|---|---|---|---|
| Age→TNFR1→COVID | 12.1 | 4.3 – 26.2 | <0.001 |
| Age→GDF15→COVID | 37.6 | 12.6 – 97.2 | 0.002 |
| TNFR1→IL-10→COVID | 5.0 | 0.9 – 36.0 | 0.036 |
| TNFR1→GDF15→COVID | 32.7 | 12.6 – 99.2 | 0.002 |
